# Working Memory Guides Perceptual Decisions Through Fast Capture and Slow Drift

**DOI:** 10.1101/2025.10.16.682875

**Authors:** Hyung-Bum Park, Weiwei Zhang

## Abstract

The top-down influence of working memory (WM) can manifest as both attentional capture and small systematic biases in perceptual judgment (i.e., “tinted lens” effect). Yet it remains unclear whether these influences arise from a single mechanism or reflect functionally distinct processes operating over different timescales. Across two experiments, we embedded a perceptual estimation task during the delay interval of a WM task and recorded time-resolved mouse trajectories during both perceptual matching and subsequent WM tests. Hierarchical Bayesian mixture modeling revealed robust bidirectional attraction between memory and perception. Time-resolved analyses of mouse trajectories further revealed two distinct components: an early, endpoint-inconsistent deviation that varied with movement onset latency, and a slower, endpoint-consistent drift that closely tracked biases in the final report. This pattern is consistent with a fast, capture-like influence of the WM template, and a sustained bias in the evolving decision, respectively. Notably, the prospective influence of WM on perception expressed both early deviation and sustained drift, whereas the retrospective influence of perception on WM primarily involved the sustained component. These findings indicate that WM shapes perceptual decisions through at least two temporally distinct contributions, and illustrate how time-resolved trajectories can reveal the dynamic structure of top-down influences within single trials.

**Statement of Relevance:** Our minds do not simply record the external world as it is, but reconstruct it actively through the lens of past experience and internal goals. Using time-resolved mouse-tracking, this study shows that working memory (WM) shapes perceptual decisions through influences that unfold over different timescales. WM can produce an early capture-like pull toward memory-matching information while also producing a slower, sustained bias aligned with the final perceptual judgment. These findings suggest that WM serves as an active and flexible source of prior information that dynamically links past and present in shaping perceptual decisions.

---

Working memory (WM) is a core cognitive system for maintaining task-relevant information and for preparing the perceptual system to interact with upcoming input (Cowan, 2001; Nobre & Stokes, 2019). Rather than functioning solely as a short-term storage, WM dynamically configures processing priorities by pre-activating feature-selective neural codes, thereby linking prior knowledge and current perceptual decisions within a predictive control framework (Myers et al., 2017; Summerfield & de Lange, 2014). In this view, WM provides an internal context that shapes how sensory evidence is sampled, weighted, and integrated over time.

This top-down influence has been studied in at least two forms. First, the features instantiated in WM can serve as an attentional template that biases selection toward matching inputs through competitive selection (Desimone & Duncan, 1995; Wolfe, 1994). Behaviorally, this often appears as attentional capture, wherein attention is automatically drawn toward WM-matching input even when such capture conflicts with current task demands (Olivers et al., 2006; Soto et al., 2008; Park & Zhang, 2024; Won & Zhang, 2025). Second, and more controversially, WM may directly alter how features appear perceptually, functioning as a “tinted lens” that pulls new sensory information toward remembered features (Teng & Kravitz, 2019; Teng et al., 2023). This tinted-lens account posits that WM content can shift nearby sensory representations toward itself, producing reliable systematic distortions in perceived color, orientation, or shape. This theoretical framework aligns with sensory-recruitment theories proposing that sensory cortices contribute simultaneously to both perception and memory maintenance (Harrison & Tong, 2009; Serences et al., 2009; Rademaker et al., 2019), as well as attractor network models in which stable memory states act as attractor basins that bias the readout of sensory evidence (Burak & Fiete, 2009; Wimmer et al., 2014). Historically, these two behavioral signatures have been investigated as largely separate phenomena using distinct methodological approaches, leaving open whether they reflect different stages of a shared process or functionally distinct computational operations that unfold across different timescales.

Answering these questions requires a method capable of tracking the evolution of WM-related influences within a single trial. Conventional measures such as accuracy and reaction time are insufficient as they cannot distinguish early transient influences from later sustained biases. Neurophysiological evidence demonstrates that WM’s influence can indeed operate on multiple timescales. For example, event-related potential studies have documented that WM-matching items modulate early visual components (e.g., P1/N1, around 100–200 ms post-stimulus), consistent with rapid attentional modulation, as well as later components associated with conflict resolution and response mapping, which occur over 500 ms post-stimulus (Luck et al., 2000; Tan et al., 2014). These findings suggest the presence of temporarily distinct contributions, yet clear behavioral evidence isolating multiple WM-driven influences within the same perceptual decision has remained elusive.

The present study employs time-resolved mouse-tracking to examine how WM shapes perceptual decisions moment-by-moment. Time-resolved cursor trajectories provide a process-level readout of evolving choices and can reveal subtle early deviations as well as slower drifts that precede the final report (Park, 2025). This approach allows us to operationally separate two trajectory components without assuming that either component uniquely identifies a single cognitive mechanism. First, we refer to an early, transient trajectory curvature toward the WM-matching feature that cannot be explained by the final endpoint response as the capture-like component (Theeuwes et al., 2005). Such curvature is consistent with WM-driven attentional guidance, but could also reflect early competition between response tendencies associated with the WM-matching and task-relevant features. Second, we refer to the sustained, endpoint-consistent deviation as the shift component, which indexes bias aligned with the final response without by itself establishing whether that bias arises at a perceptual, decisional, or response-readout stage. To accomplish this, we adopted an analytic method to decompose the overall trajectory into two components, an early endpoint-inconsistent deviation and a sustained endpoint-consistent deviation (Park, 2025; see *Data Analysis* for detail).

Across two experiments, participants memorized a briefly presented sample color, performed a perceptual matching task embedded during the memory delay, and then completed a WM test. Experiment 1 examined how WM influenced ongoing perceptual decisions, and Experiment 2 examined the reciprocal influence of perceptual decisions and subsequent WM recall to investigate how the priority state of the maintained representation (i.e., prospective vs. retrospective influence) relates to early-versus-late components of the decision process. Across both experiments, we show that WM content produces an early capture-like influence and a sustained endpoint-consistent bias that emerge on distinct temporal scales. Specifically, the capture-like component emerges and dissipates rapidly, consistent with transient priority-based competition, whereas the endpoint-consistent bias develops gradually and persists throughout movement, consistent with a sustained integrative influence on the evolving decision. Together, these findings reveal how temporally distinct WM-related influences jointly shape evolving perceptual decisions.

## General Method

### Participants

Thirty-three (19 female) and twenty-one (14 female) college students at the University of California, Riverside participated in Experiment 1 and 2, respectively, in exchange for course credit. Participants were aged between 18 and 24 years, reported normal or corrected-to-normal vision and normal color vision. Informed consent was obtained prior to participation, and all experimental protocols were approved by the University of California, Riverside Institutional Review Board.

For Experiment 1, the target sample size was determined based on previous work adopting a closely related experimental and mouse-tracking procedure (Park & Zhang, 2024). A priori power analysis in G*Power 3.1 (Faul et al., 2009) for a two-tailed paired *t*-test with *α* = .05 and a predicted effect size of *dz* = 0.50 indicated that 34 participants would provide approximately 80% power. The sample size for Experiment 2 was guided by the robust within-subject bias observed in Experiment 1 (*d* = 1.29), thus moderate *N*s are sufficient to detect reliable within-subject attraction effects. It should be noted that the primary statistical inferences in both experiments were based on hierarchical Bayesian modeling and resampling-based permutation tests (see *Data Analysis* for detail). The Bayesian analyses yielded decisive posterior credible intervals for the population-level parameter of key interest, and the permutation tests detected the predicted dissociation effects, supporting the adequacy of the sample size.

## Stimuli and Procedure

Stimuli were presented on a 60 Hz LCD monitor calibrated with an X-Rite i1Pro spectrophotometer, with a grey background (15.1 cd/m²) at a viewing distance of 60 cm. Task procedures were identical in Experiments 1 and 2, except for a test array (Figure 1). Each trial began with central fixation (800 ms), followed by a sample display (100 ms) of a single colored square (1.6° × 1.6°), which appeared at one of four equally spaced locations on an imaginary circle (radius 4.3°) centered at fixation. The memory color was selected randomly from 180 evenly spaced hues on a circular CIELAB color space (Zhang & Luck, 2008).

**Figure 1.**
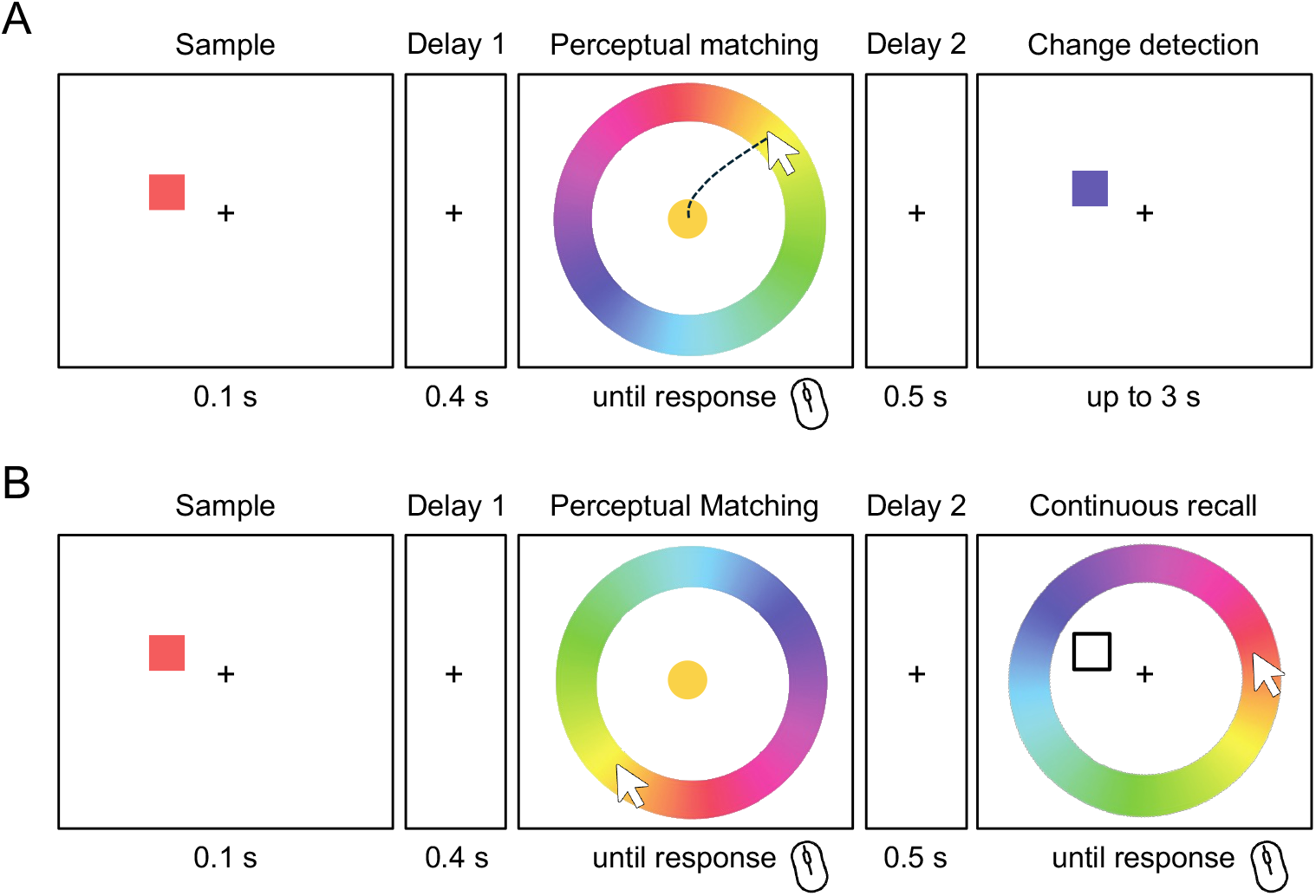
Task procedure. (A) Experiment 1 combined a perceptual matching task with a subsequent working memory (WM) change-detection task. After viewing a sample color and a short delay, participants reported the color of a continuously visible target using a color wheel. Mouse trajectories were recorded from movement onset to the selected color, as illustrated by the example path. This perceptual-matching response was followed by a second delay and a change-detection judgment, for which change magnitude varied in 60° steps in both directions (−180°:60°:180°). (B) Experiment 2 combined perceptual matching with continuous WM recall. After the sample and delay, participants performed the same perceptual-matching response with mouse-trajectory recording, followed by a second delay and continuous recall of the memorized color using a color wheel. Mouse trajectories were also recorded during the WM recall response.

After an unfilled 400-ms delay, the perceptual matching display appeared. A colored disk (1.6° diameter) replaced the central fixation point, surrounded by a continuous color-wheel (radius 8.2°, thickness 2.2°) with 180 evenly distributed hues from the same color space. The target color for the perceptual matching task was offset by either 40° clockwise (CW) or counterclockwise (CCW) from the memorized color in the sample display. This relative offset was strategically chosen to fall within the similarity range that maximizes attraction in previous serial-dependence research (Fischer & Whitney, 2014). The target item remained visible until response, and participants reported the perceived hue of the target by moving the mouse cursor from the center of the screen to the best-matching color on the color-wheel with a click. Mouse trajectories (x and y coordinates and time relative to the cursor onset) were recorded continuously. Immediately after the click, 500 ms visual feedback appeared on the color-wheel, with a cross marking the color the participants picked and an arrow marking the target color. The color-wheel was randomly rotated on every trial to prevent fixed spatial mapping of the colors to locations on the wheel. The mouse cursor was re-positioned at fixation after the visual feedback.

Following another unfilled 500 ms delay, a test display occurred, which differed by experiment. Experiment 1 adopted a change-detection task, in which the test array presented a single probe at the original memory item location, and participants indicated whether the probe matched or mismatched the remembered color by pressing one of two shoulder buttons on a gamepad (e.g., L1 for “no-change”, L2 for “change”). The participants used their dominant hand (right hand for all subjects) for the mouse clicking for the perceptual matching task and the nondominant hand for the change-detection task. Change and no-change trials occurred with equal probability. On change trials, the probe color rotated 60°, 120°, or 180° in either direction from the memory color with equal probability. Because the preceding perceptual matching target was always positioned 40° CW or CCW relative to the memory color, this systematic manipulation of change magnitude allowed us to test whether the perceptual judgment made during WM maintenance influenced the subsequent representation of the memory item. Auditory feedback followed each response, consisting of a 500 ms low-pitch tone for correct answers and a high-pitch tone for errors.

Experiment 2 adopted a continuous recall paradigm, in which the test display consisted of an outlined square at the original location of the memory item and another color-wheel that had a circular rotation to random degrees in order to prevent a strategic mapping between the stimulus and color-wheel location during the previous perceptual matching task. Participants reproduced the remembered color by moving the mouse cursor away from the center of the screen and then clicking the best-matching color on the wheel. Immediate feedback was provided in the same manner as that for the perceptual matching task. In both experiments, the mouse cursor was automatically re-centered at fixation after each click, and all participants completed 300 trials in total, divided into three blocks with brief breaks.

## Data Analysis

### Data and code availability

All primary and processed data, as well as custom analysis codes, are publicly available via the Open Science Framework repository (https://osf.io/3pyjh).

### Hierarchical Bayesian Mixture Modeling

Perceptual matching error in Experiment 1 and 2 was defined as the signed angular distance in degrees between each reported color and the corresponding physical target color in the perceptual matching task. Similarly, memory recall error was defined as the signed angular distance in degrees between the reported and the original memory colors during WM recall in Experiment 2. These errors were modeled with a hierarchical Bayesian mixture model that extends the standard continuous estimation framework (Zhang & Luck, 2008). The model is expressed as a mixture of two distributions representing whether recall responses are based on a noisy target representation or non-informative guesses (Figure 2). Specifically, the von Mises component captures responses based on noisy feature representations with mean (*μ*) that indexes inaccuracy or bias and concentration *κ* that indexes precision (reported as imprecision *σ* converted from *κ*). The uniform component captures guesses or lapses with mixture weight *g*, the height of the uniform probability distribution. Although guessing was expected to be nearly minimal in the perceptual matching task since the target remained visible until response, the same modeling approach was applied to both perceptual matching and continuous WM recall errors in Experiment 2. This modeling approach permits a principled cross-task comparison of bias magnitude while accounting for occasional lapses or memory failures.

**Figure 2.**
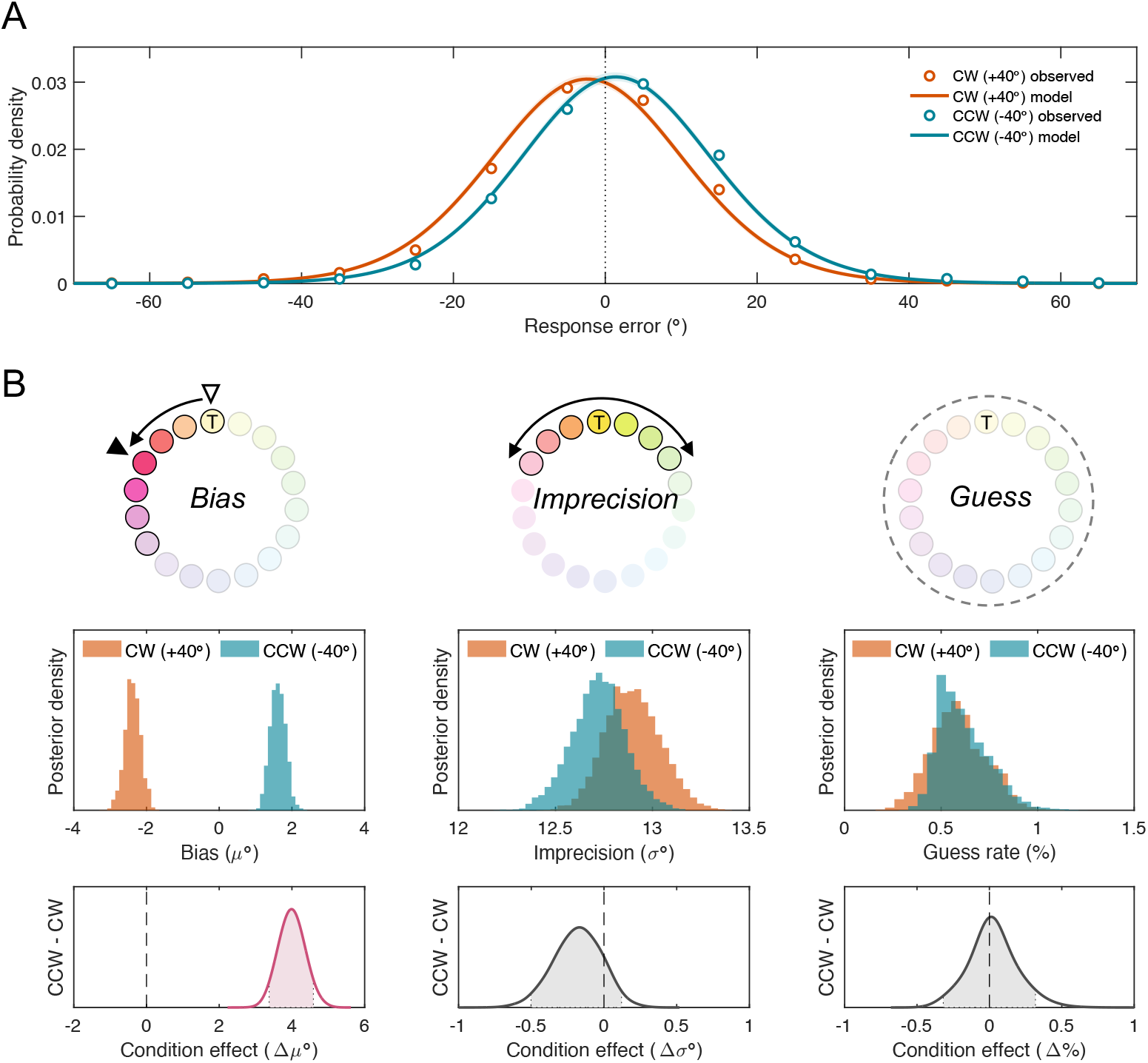
**Hierarchical Bayesian parameter estimates for Experiment 1**. (A) Observed error distributions and posterior Mixture model fits for the CW (+40°; orange) and CCW (−40°; teal) conditions. (B) Posterior distributions of the population-level parameters for bias (*μ*), imprecision (*σ*), and guess rate (*g*). The bottom panels show posterior distributions of the condition effects (CCW − CW). A credible condition difference was observed for bias (Δ*μ*), with 95% highest density intervals (HDI95%) did not include zero. Shaded areas represent HDI95%.

The bias effects in serial-dependence and WM are typically reliable but numerically small, often about a few degrees (Barbosa & Compte, 2020; Park, 2025). Hierarchical Bayesian modeling offers key advantages for this context, because it simultaneously accounts for multiple sources of variability (e.g., across participants, conditions, and trials), thereby stabilizing estimation of small effects (Oberauer et al., 2017; see also Frischkorn & Popov, 2025). This approach allows noisy parameter estimates to be shrunk toward the population mean, leading to more robust and reliable results (Shiffrin et al., 2008). The choice of reasonably informative to non-informative priors was determined following previous studies with a similar modeling approach (Oberauer et al., 2017; Park & Zhang, 2024). We took a total of 4,000 samples after 4,000 warm-ups with four chains of Markov chain Monte Carlo sampling. It yielded posterior matrices of population-level parameters sized, for example, 12,000 (samples) × 33 (subjects) × 2 (conditions) for Experiment 1. Convergence was assessed using the Gelman-Rubin R^ statistic, with all population-level parameters showing values equal or close to 1.0 (Gelman & Rubin, 1992). Statistical inferences are reported in terms of posterior means and the 95% highest density credible intervals (HDI_95%_; Kruschke, 2015).

### Mouse Trajectory

Mouse cursor trajectories were sampled throughout each response from stimulus onset until the click on the color-wheel during both the perceptual matching task (Experiments 1 and 2) and the subsequent WM recall task (Experiment 2). Movement onset was defined as the first sample in which the cursor moved more than five pixels from the central starting position. To compare trajectories across trials and participants, each trajectory was time-normalized from movement onset (0%) to the final response (100%). On average, participants initiated the cursor movement at about 25.7% [*CI_95%_*: 22.6%, 28.8%] of the normalized time scale.

To place all trials in a common spatial reference frame, each trajectory was rotated so that the true target color mapped to the top of the response color-wheel with Cartesian coordinate (0, *r*), where *r* is the wheel radius. This normalization ensured that any horizontal deviation directly reflects attraction toward or repulsion away from the WM-matching color during perceptual matching, or relative to the intervening perceptual target during WM recall.

The direction and magnitude of trajectory curvature were summarized as the signed area under the curve (AUC), computed as the integral of horizontal cursor offsets across the normalized time axis. After confirming that there was no reliable difference in curvature magnitude between trials where the perceptual target lay clockwise (CW) or counterclockwise (CCW) of the WM item, *t*(32) = 0.08, *p* = .933, Cohen’s *d* = 0.01, we flipped the sign for the CW trials. Consequently, positive AUC values for perceptual matching indicate attraction toward the WM representation, and negative AUCs for WM recall in Experiment 2 indicate attraction toward the earlier perceptual decision.

To distinguish transient attentional capture-like component from the more sustained, endpoint-consistent bias, we employed a decomposition method utilizing an endpoint-bias correction (Park & Zhang, 2022). For each trial, we constructed a reference path connecting the starting point at fixation to the final click location while preserving the same radial progress over time as the observed trajectory. This reference path captures the portion of the movement that is expected solely from the final response location. Thus, an early deviation toward the WM color that is later corrected remains in the capture component, whereas displacement that persists in the final response contributes to the shift component. Subtracting this reference path from the overall trajectory therefore yielded trajectory deviations for the *capture* component that cannot be explained by the endpoint. The complementary *shift* component was defined as the residual (i.e., overall trajectory = capture + shift) and indexes the portion of curvature that is geometrically consistent with the endpoint displacement. The overall AUC is thus the sum of capture (AUC*_capture_*) and shift (AUC*_shift_*).

Because AUC*_shift_*, by construction, reflects curvature aligned with the chosen endpoint, its magnitude is expected to covary with the final response bias. Accordingly, we do not treat AUC*_shift_* as a statistically independent source of evidence about the final report. Instead, the decomposition serves as a variance-partitioning tool that provides a principled way to test whether different portions of the trajectory associate with theoretically relevant predictors, such as movement initiation latency or final judgment bias, in ways that could reveal functional differentiation among WM’s top-down influences. This approach mirrors prior decomposition strategies in serial dependence and dynamic decision-making research, where early deviations and late drifts reflect separable computational influences on evolving choices, even in the opposite directions (e.g., early repulsion vs. later attraction; Park, 2025).

Beyond area-based point estimates, we further conducted time-resolved analysis using a recently developed destination-vector transformation (DVT) to capture moment-by-moment dynamics of decision making (Park & Zhang, 2024; Figure 5A). Rather than focusing on the instantaneous cursor position, DVT estimates the projected location on the continuous response scale at each time point, based on the direction of the local movement vector connecting two consecutive samples. For a trajectory with *n* samples, DVT produces *n−1* destination-vectors in circular degrees, providing a time series of “intended destinations”. We applied DVT separately to the overall, capture, and shift trajectories, and averaged the resulting time series across trials and participants to measure the temporal dynamics of how WM or recent perceptual input shaped the evolving response tendencies. Time-resolved differences between conditions and components were evaluated using within-subject cluster-based permutation tests (10,000 label swaps), providing a robust assessment of temporal intervals in which the components diverged.

## Results

### Experiment 1: Perceptual Matching

#### Endpoint Bias Toward Working Memory

Perceptual matching errors as the endpoint of the mouse trajectory showed the predicted attraction toward the WM-matching color on the response color-wheel. Mean circular error was negative in the CW condition (i.e., the perceptual target color is CW to the memory item; *M* = −2.44° [*CI_95%_*: −3.03°, −1.85°]) and positive in the CCW condition (+1.89° [*CI_95%_*: +1.09°, +2.68°]), yielding a reliable difference between conditions, *t*(32) = −7.31, *p* < .001, Cohen’s *d* = −1.29. A Hierarchical Bayesian Mixture model confirmed these attraction effects (Figure 2). Posterior distributions of the population-level bias (*μ*) parameter were in opposite directions for CW and CCW trials, and the 95% HDI for their contrast excluded zero, Δμ_CCW−CW_ = +4.00° [HDI_95%_: +3.37°, +4.59°], indicating robust attraction toward the WM color. Imprecision (*σ*) and guess rate (*g*) did not differ credibly between CW and CCW trials (Δ*σ =* 0.18° [HDI_95%_: −0.12°, 0.50°]; Δ*g* = −0.03% [HDI_95%_: −0.32%, 0.32%]).

#### Mouse Trajectory Deviation Toward Working Memory

Next we assessed the entire mouse trajectory beyond the endpoint to reveal the temporal dynamics of the decision process. The overall trajectory for both CW and CCW trials showed a clear early deviation toward the WM color, which peaked soon after movement onset and declined as the movement progressed (Figure 3), manifested as the sizable horizontal deviations and the AUC*_overall_* (13.8 px [*CI_95%_*: 10.9 px, 16.8 px]), *t*(32) = 9.28, *p* < .001, Cohen’s *d* = 1.64). Moreover, AUC*_overall_* scaled with how fast participants initiated mouse cursor movement upon perceptual target onset. A within-subject median-split on movement onset latency revealed larger deviations on early-onset trials (AUC*_early_* = 24.1 px [CI_95%_: 20.0 px, 28.2 px]), than on late-onset trials (AUC*_late_* = 9.6 px [CI_95%_: 6.8 px, 12.3 px]), *t*(32) = 8.05, *p* < .001, Cohen’s *d* = 1.42. This speed contingency suggests that rapid movements are subject to a brief capture-like influence toward the WM-matching item, thereby magnifying overall trajectory curvature. In contrast, movement onset latency did not reliably influence the final perceptual error. Biases were similar for early-onset trials (2.32° [CI_95%_: 1.64°, 3.00°] and late-onset trials (1.88° [CI_95%_: 1.21°, 2.55°]), *t*(32) = 1.23, *p* = .229, Cohen’s *d* = 0.22. The absence of a reliable endpoint difference argues against a simple account in which the endpoint bias merely reflects residual under-correction of the initial deviation, and instead supports a functional dissociation between early trajectory curvature and the sustained endpoint bias. Because AUC*_overall_* conflates these influences, we decomposed the measure to isolate endpoint-inconsistent and endpoint-consistent contributions to the evolving response.

**Figure 3.**
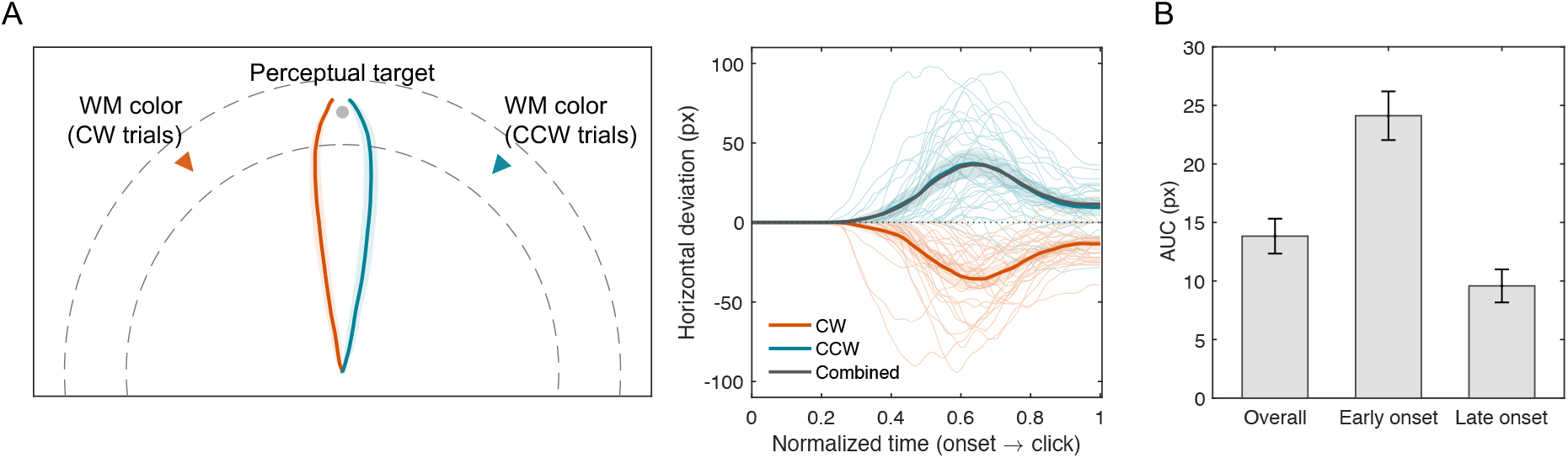
Mouse-tracking evidence for WM-induced perceptual bias in Experiment 1. (A) Average trajectory paths on a schematic display (left) and time-normalized horizontal trajectory deviations (right), plotted separately for CW (+40°) and CCW (−40°) conditions. Thin lines show individual participant means, thick lines show group means, and shaded bands indicate 95% confidence intervals. The gray line shows the combined trajectory deviation after aligning CW and CCW trials to a common direction. (B) Area-under-the-curve (AUC) of trajectory curvature for all trials and for early- and late-onset trials defined by a within-participant median split of movement onset latency. Error bars indicate standard errors of the mean.

#### Functional Dissociation: Capture versus Shift

To determine whether earlier, endpoint-inconsistent deviations and later, endpoint-consistent drifts show different functional relationships with behavioral predictors, overall curvature was decomposed with an endpoint-bias correction (Figure 4A). For each trial, a reference line connecting the start and endpoint was subtracted from the observed path, yielding the *capture* component (deviations not explained by the endpoint). The *shift* component was defined as the residual (i.e., overall = capture + shift). We computed AUC for each component and conducted a set of individual-differences tests relating these components to movement onset latency and to the magnitude of the final perceptual shift.

**Figure 4.**
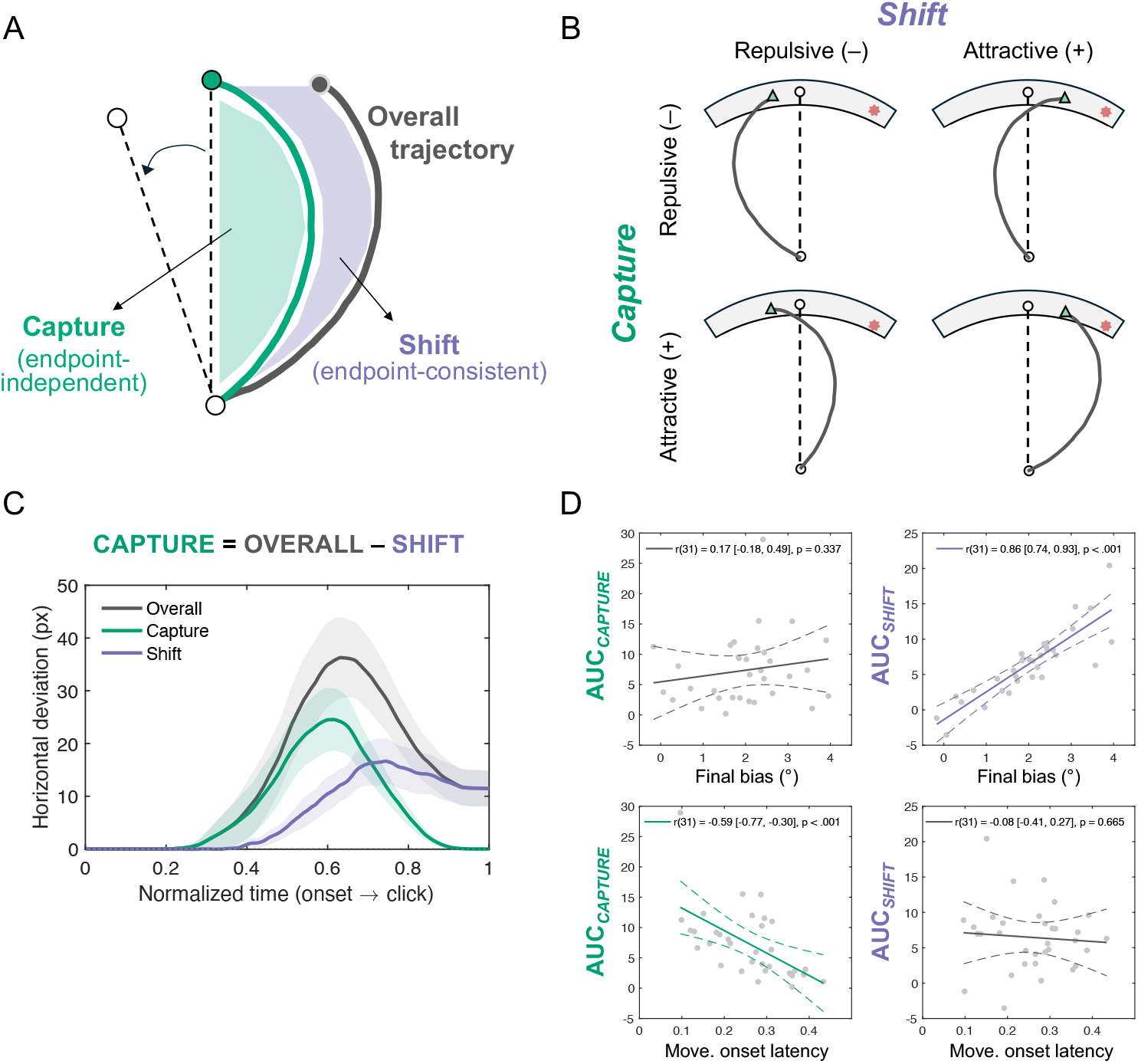
Decomposition of trajectory bias into capture and shift components in Experiment 1. (A) Schematic illustration of the endpoint-based decomposition. The observed overall trajectory (gray) was decomposed into an endpoint-inconsistent capture component (green) and an endpoint-consistent shift component (purple). The capture component was obtained by subtracting the endpoint-consistent reference path from the observed trajectory (B) Conceptual 2 × 2 examples showing that capture and shift can independently take attractive (positive) or repulsive (negative) deviations from the source of bias (red markers). White circles and green triangles indicate the true target location and the judgment endpoint, respectively. (C) Average time course of horizontal deviation for the overall, capture, and shift trajectories, time-normalized from movement onset to the final click. Shaded bands indicate 95% confidence intervals (CI95%). (D) Correlations of area-under-the-curve (AUC) measures for capture and shift with final perceptual bias (top row) and movement onset latency (bottom row). Solid lines show the best linear fit and dashed lines the CI95%. Regression lines are shown in the corresponding component color for statistically significant effects and in gray for nonsignificant effects.

This decomposition revealed distinct prediction patterns across the two components (Figure 4D). Movement onset latency reliably predicted AUC*_capture_*, *r*(31) = −.59 [*CI_95%_*: −.77, −.30], *p* < .001, such that faster initiations were associated with larger endpoint-inconsistent deviations, whereas it showed little relationship with AUC*_shift_*, *r*(31) = −.08 [*CI_95%_*: −.41, .27], *p* = .654. In contrast, the magnitude of the final endpoint bias was strongly associated with AUC*_shift_*, as expected given that it is defined with respect to the endpoint path, *r*(31) = .86 [*CI_95%_*: .74, .93], *p* < .001, but was weakly related to the AUC*_capture_*, *r*(31) = .17 [*CI_95%_*: −.18, .49], *p* = .336. Once endpoint-consistent curvature was factored out, the remaining capture-only component showed a selective relationship with movement onset latency and minimal association with the final perceptual bias. This pattern suggests that AUC*_capture_* tracks an early, initiation-linked influence of working memory that is functionally distinct from the endpoint-consistent bias reflected in AUC*_shift_*.

#### Time-Resolved Decision Dynamics: Destination-Vector Series

The analyses of movement latency and component dissociation suggested that WM influences manifest as both a transient attention capture-like influence and a sustained endpoint-consistent bias. To track how these influences unfolded over time, we used DVT to reconstruct evolving destination estimates throughout the movement trajectory (Figure 5A). The decomposition (Figure 5B) mirrored the AUC findings. The capture time series (DV_CAPTURE_) rose sharply at movement onset and returned toward baseline before the response was completed, consistent with a transient attentional capture. The shift component (DV_SHIFT_) remained positive throughout the trajectory, emerging early and persisting until the response, consistent with a steady representational shift.

To directly compare these temporal dynamics, we examined slope differences between the capture and shift effects. Cluster-based permutation testing (10,000 iterations) revealed an early interval (5% to 15% normalized time) where capture increased more steeply than shift (i.e., capture slope > shift slope, *p*_cluster_ < .05), followed by a late period (25% to 95% normalized time) where the difference reversed in sign (i.e., capture slope < shift slope, *p*_cluster_ < .05; Figure 5C). These results indicate that WM exerts dissociable influences on decision dynamics. A rapid capture-like contribution dominates early in the movement, whereas a slower, sustained shift persists throughout and aligns with the endpoint bias.

Building on the selective relationship between movement onset latency and AUC*_capture_* shown in Figure 4D, we next asked whether this relationship was also evident in the time course of the DVT-derived capture and shift components. Trials were divided within each participant using a median split of movement onset latency (Figure 6). Mean movement trajectories showed greater WM-directed deviation on early-onset than late-onset trials (Figure 6A). Consistent with the AUC analysis, the capture component was substantially larger on early-onset trials, with a significant difference spanning 15.8% to 89.5% of normalized DVT time (*p*_cluster_ < .001; Figure 6B). The shift component was also numerically larger on early-onset trials, but this difference did not survive cluster correction (smallest *p*_cluster_ = .091; Figure 6C). Together, these results indicate that the relationship between earlier movement initiation and greater trajectory deviation was expressed primarily in the capture component, with no reliable time-resolved modulation of the sustained shift component.

**Figure 5.**
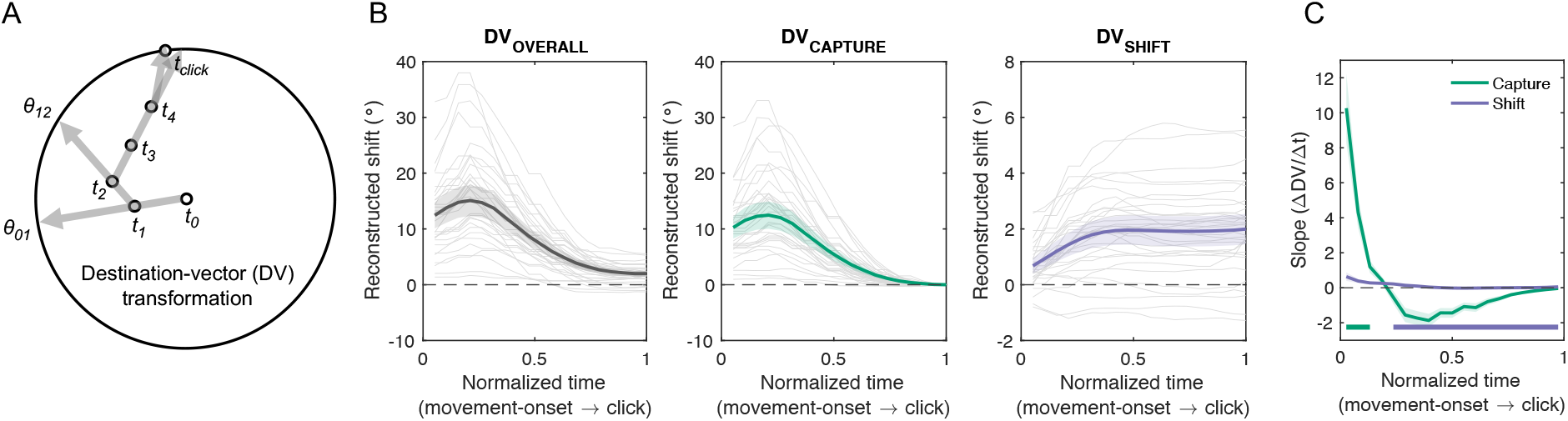
Destination-vector transformation and decomposition analyses in Experiment 1. (A) Schematic illustration of the destination-vector (DV) transformation method, which represents each movement segment as an angle relative to the target based on two consecutive moments, allowing reconstruction of intended feature values across time (Park & Zhang, 2024). (B) Reconstructed DV dynamics for overall deviation (gray), capture (green), and shift (purple), from left to right. Thin gray lines show individual participant DV series and thick lines with shaded bands indicate group means and 95% confidence intervals. (C) Green and purple horizontal bars denote time windows in which capture and shift, respectively, showed the steeper slope in significant cluster-based permutation effects (*p* < .05).

**Figure 6.**
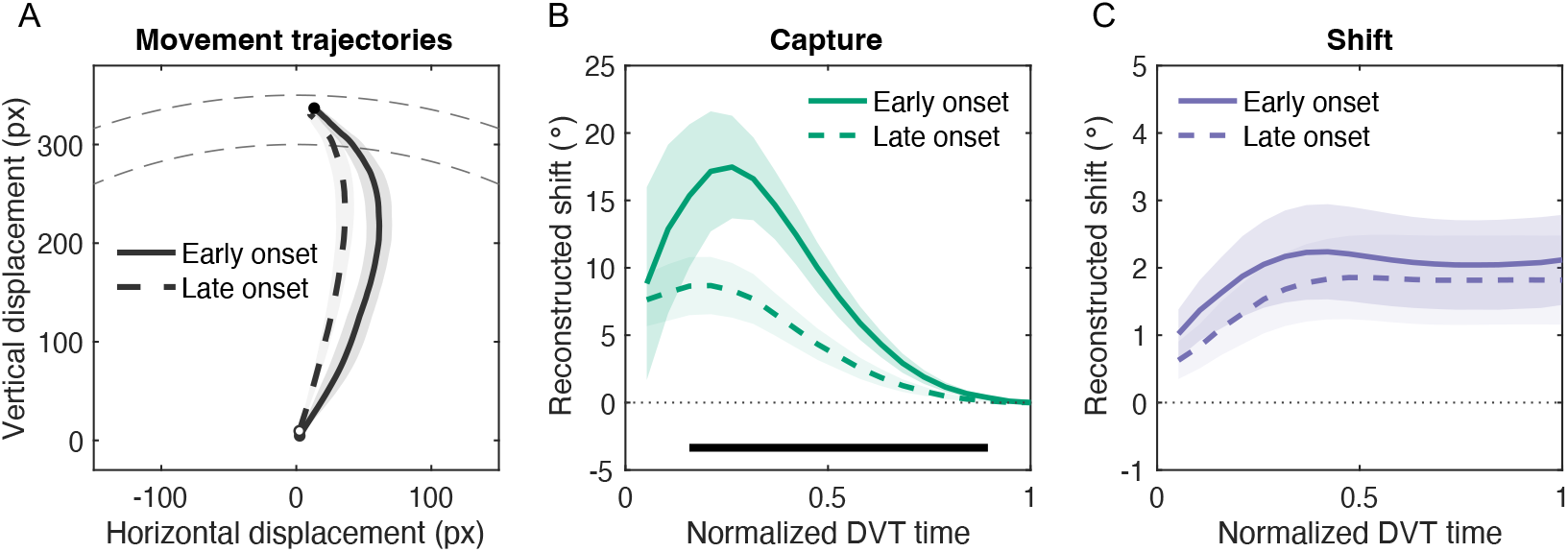
Movement-onset latency modulates time-resolved trajectory dynamics in Experiment 1. (A) Mean movement trajectories for early- and late-onset trials, defined by a within-participant median split of movement-onset latency. Shaded regions indicate 95% confidence intervals for horizontal displacement across participants. Dashed semicircles provide the display reference frame. (B) Destination-vector time course of the capture component for early- and late-onset trials. The black horizontal bar marks the significant early-versus-late cluster identified by cluster-based permutation testing. (C) Destination-vector time course of the shift component for early- and late-onset trials. Shaded regions indicate 95% confidence intervals.

## Experiment 1: Working Memory Change Detection

The WM change-detection task involved systematic manipulations of change magnitude, varying memory-to-probe color offset every 60° steps. To test whether perceptual matching influenced WM content retrospectively, we modeled the psychometric function relating the probability of “same” responses as a function of change magnitude, where 0° represents no-change trials (Figure 7A). The group-aggregated curves were fit with a Gaussian function to estimate the point of subjective equality (PSE, *μ*_Gauss_), slope (*σ*), amplitude (*Amp*), and *lapse* rate, separately for CW and CCW curves. This revealed a reliable shift of the PSE (*μ*_Gauss_) between CW (+2.59°) and CCW (−6.48°) conditions. Bootstrap distributions of the PSE estimates are shown in Figure 7B. For statistical inference, subject-level bootstrap resampling (10,000 iterations) provided CI_95%_ and two-sided *p* values for the differences in these estimates across the conditions. The PSE (*μ*_Gauss_) showed a reliable condition difference between the CW and CCW conditions (9.11° [CI_95%_: 4.67°, 13.42°], *p*_boot_ < .001). None of the other parameters showed reliable differences (for *Amp*, −0.01 [CI_95%_: -0.03, 0.02], *p*_boot_ = .529; for *σ*, −1.25° [CI_95%_: −4.12°, 1.68°], *p*_boot_ = .384; for *lapse*, −0.012 [CI_95%_: −0.026, 0.001], *p*_boot_ = .071).

**Figure 7.**
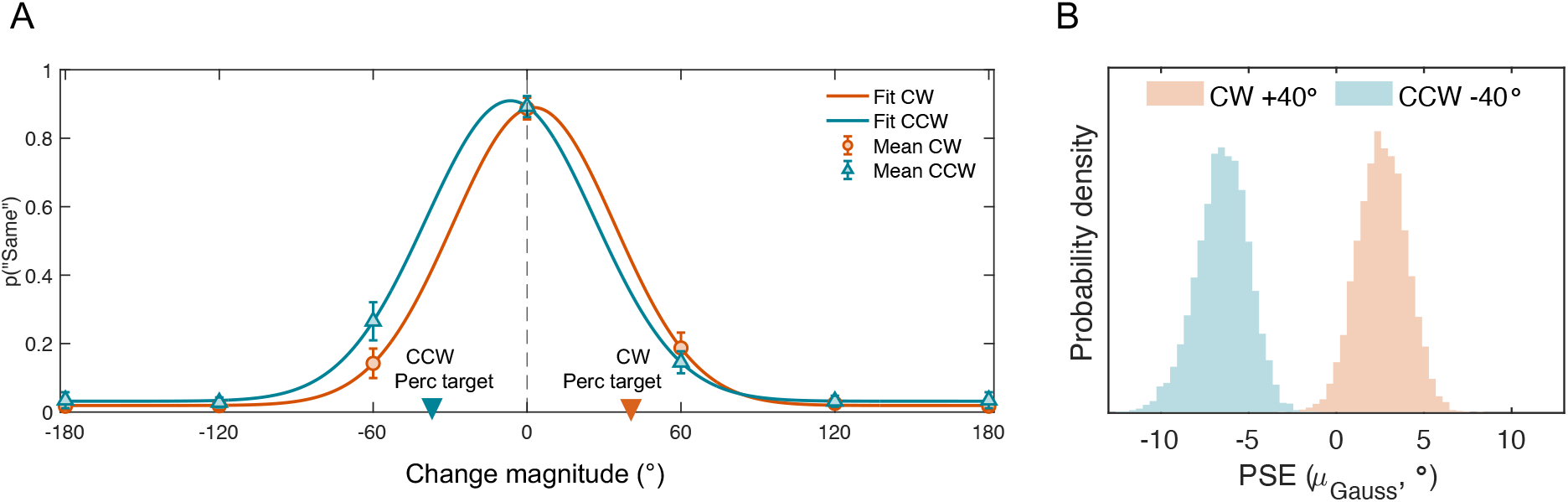
Working memory change-detection performance in Experiment 1. (A) Proportion of “same” responses as a function of change magnitude, with Gaussian fit curves (solid lines) shown separately for CW (+40°) and CCW (−40°) trials. Markers with error bars indicate group means and 95% confidence intervals. Inverted triangles along the x-axis denote the corresponding locations of the preceded perceptual target features. (B) Bootstrap distributions of the point of subjective equality (PSE, *μ*Gauss) estimated separately for CW and CCW conditions.

The change-detection results demonstrate that WM judgments are reliably biased in the direction of the preceding perceptual decision. Experiment 2 replaced a binary change-detection with a continuous WM recall procedure following the perceptual matching task (Figure 1B), allowing a direct comparison between attraction observed during perceptual matching and that during mnemonic recall.

## Experiment 2

### Endpoint Bias between Perceptual Matching and Working Memory

Hierarchical Bayesian Mixture modeling of continuous-report errors provided converging evidence for reciprocal attraction between WM and perception (Figure 8). Posterior predictive fits closely tracked the observed error distributions in both phases (Figure 8A). Posterior distributions of the *μ* parameter revealed opposite biases across the responses (Figure 8B). During perceptual matching, μ shifted toward the WM color, replicating Experiment 1 (Δμ_CCW−CW_ = 3.06° [HDI_95%_: 2.29°, 3.94°]). Critically, during subsequent WM recall, *μ* shifted in the opposite direction, toward the just-seen perceptual target (Δμ_CCW−CW_ = −3.63° [HDI_95%_: −5.06°, −2.34°]). Although opposite in sign, their absolute magnitudes were numerically similar, indicating reciprocal influences in the two directions. By contrast, imprecision (*σ*) did not differ credibly between CW and CCW trials in either phase (perceptual matching: Δ*σ =* −0.10° [HDI_95%_: −0.63°, 0.47°]; WM recall: Δ*σ =* −0.30° [HDI_95%_: −1.46°, 0.92°]). Guess rate (*g*) also remained low and stable (perceptual matching: Δ*g* = −0.27% [HDI_95%_: −1.27%, 0.79%]; WM recall: Δ*g* = 1.20% [HDI_95%_: −1.11%, 3.55%]). These results show that WM and perceptual judgments exerted reciprocal biases in opposite directions, with numerically similar effect magnitudes, extending the WM-to-perception effect in Experiment 1 to the complementary perception-to-WM influence.

**Figure 8.**
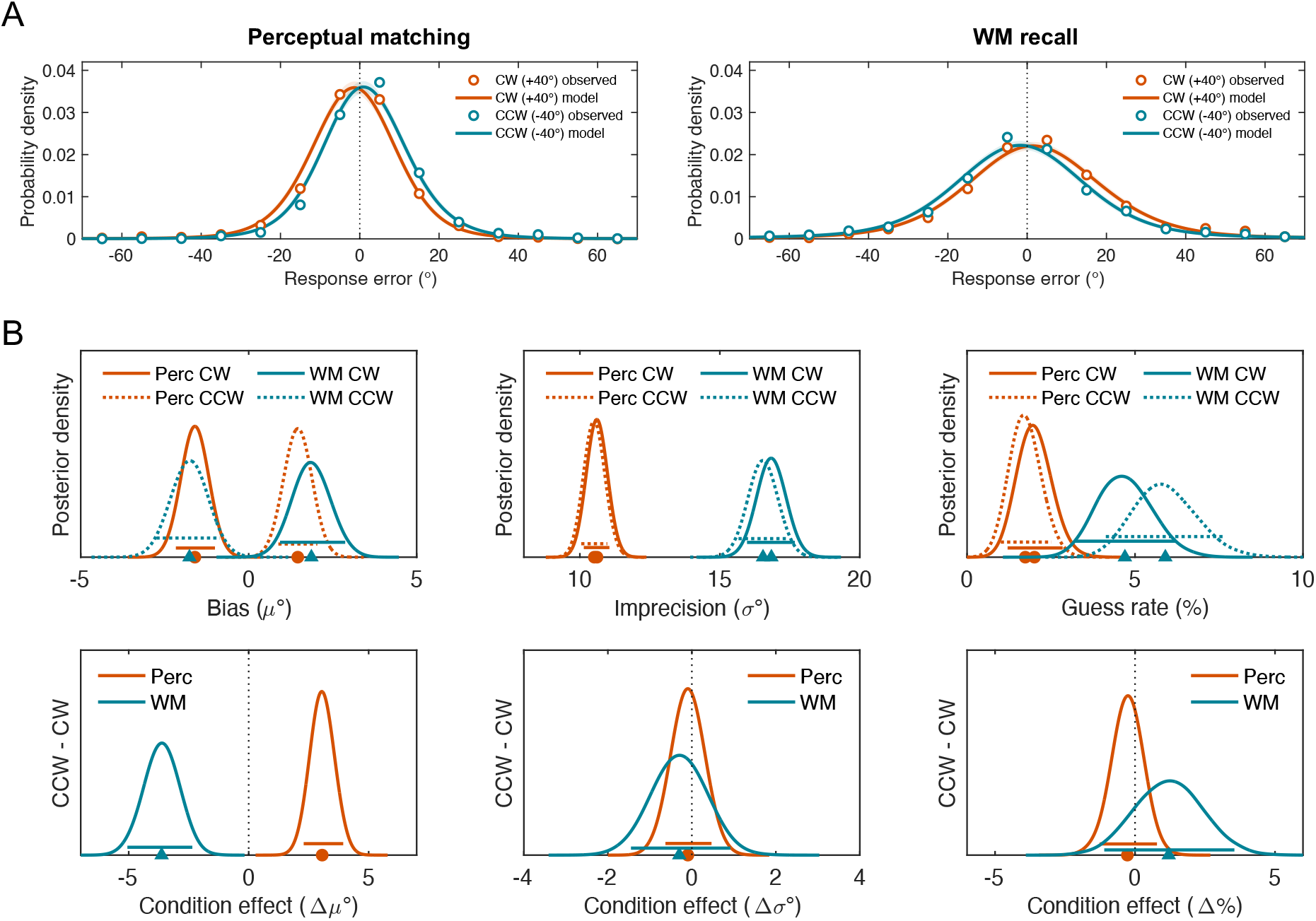
Hierarchical Bayesian Mixture modeling of continuous-report errors in Experiment 2. (A) Observed response-error distributions (open circles) and posterior predictive model fits (solid lines) for perceptual matching (left) and WM recall (right), shown separately for clockwise trials (CW +40°) and counterclockwise trials (CCW −40°). (B) Posterior distributions of population-level parameters for bias (*μ*), imprecision (*σ*), and guess rate (*g*), shown separately for perceptual matching (Perc; orange) and WM recall (WM; teal), with solid lines indicating CW and dotted lines indicating CCW. The bottom row shows posterior distributions of condition effects (CCW − CW) for each parameter. Markers and horizontal bars along the x-axis indicate posterior means and 95% highest density intervals, respectively.

### Time-Resolved Decision Dynamics: Destination-Vector Series

To examine how bidirectional biases unfolded during movement, we applied the DVT to mouse trajectories for perceptual matching and WM recall, and decomposed them into the capture (DV_CAPTURE_) and shift (DV_SHIFT_) components that showed dissociable temporal dynamics for the two responses (Figure 9). Perceptual matching showed strong early deviations toward the WM color, whereas WM recall showed a much smaller, oppositely directed early deviation. This asymmetry was primarily expressed in the capture component (DV_CAPTURE_).

**Figure 9.**
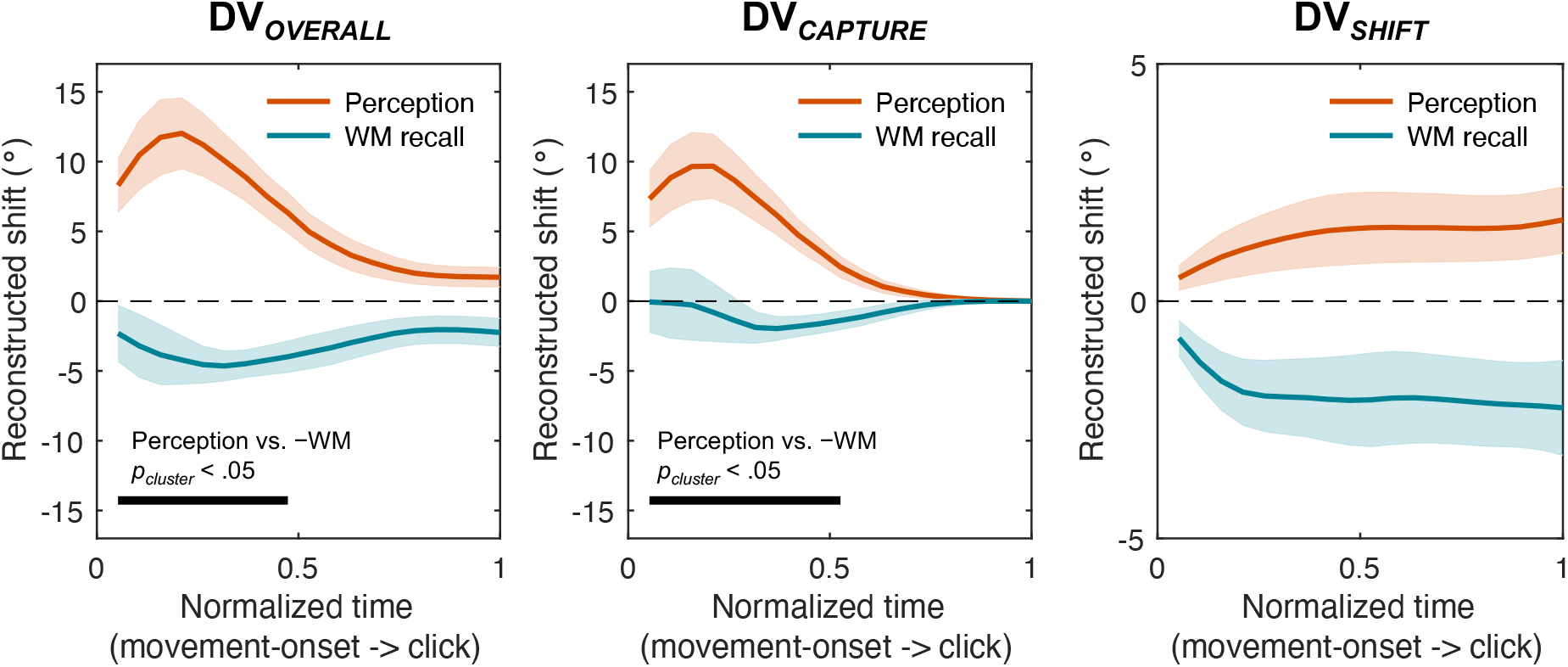
Reconstructed destination-vector (DV) dynamics and decomposition analyses in Experiment 2. Mean DV series with 95% confidence intervals over normalized time for perceptual matching (Perception; orange) and working memory recall (teal), shown separately for DV*OVERALL*, DV*CAPTURE*, and DV*SHIFT* (left to right). Black horizontal bars indicate time intervals with significant cluster-based permutation effects contrasting perceptual matching and sign-flipped WM recall (*p* < .05).

During perceptual matching, capture rose steeply after movement onset and declined thereafter, whereas WM recall showed a weaker, oppositely directed capture effect. In contrast, the shift component (DV_SHIFT_) showed steadily increasing amplitude in opposite directions for the two responses, mirroring the endpoint bias. Note, the gradual increase in DV_SHIFT_ during WM recall should not be interpreted as indicating that the mnemonic representation itself began to shift only after movement onset. Because DVT indexes the evolving destination expressed in the movement trajectory, rather than the mnemonic state before movement, a bias that was already present before movement could become progressively expressed as the recall response unfolded.

Cluster-based permutation tests (10,000 label swaps within subject) on the reconstructed destination vectors confirmed this dissociation. The capture component differed reliably between perceptual matching and WM recall from 10% to 55% of normalized movement time (*p*_cluster_ < .001), indicating stronger capture during perceptual matching. Capture was also reliably different from zero within each phase. In perceptual matching, capture differed from zero from 10% to 80% of normalized movement time (*p*_cluster_ < .001), whereas the smaller, oppositely directed capture effect during WM recall differed from zero from 35% to 65% (*p*_cluster_ = .001). For shift, no between-phase clusters were identified at the cluster-forming threshold, providing no evidence for a reliable difference in the sustained shift between phases.

Together, these results demonstrate reciprocal but asymmetrical interactions between perceptual matching and WM recall. Specifically, perceptual matching showed a substantially stronger capture component, whereas both responses exhibited sustained shifts in opposite directions. WM recall therefore showed a reliable but weaker capture-like influence from the preceding perceptual input rather than an absence of capture. This asymmetry is consistent with stronger expression of the capture component when WM contents are prospectively prioritized for ongoing behavior.

## General Discussion

The present study examined how WM shapes perceptual judgments in relation to its established role in guiding attentional selection (see Table 1). Across two experiments, we observed bidirectional attraction effects between WM and perception. Perceptual judgments were systematically biased toward the feature value held in WM, and subsequent WM recall showed bias toward the preceding perceptual decision. These findings provide time-resolved behavioral evidence that WM’s top-down influence is not well captured by a single, unitary process. Instead, the dynamics of mouse trajectories suggest that at least two temporally dissociable components contribute to the observed biases. Specifically, an early, transient influence emerges shortly after movement initiation, whereas a later, more sustained influence persists throughout the perceptual judgment.

**Table 1:**
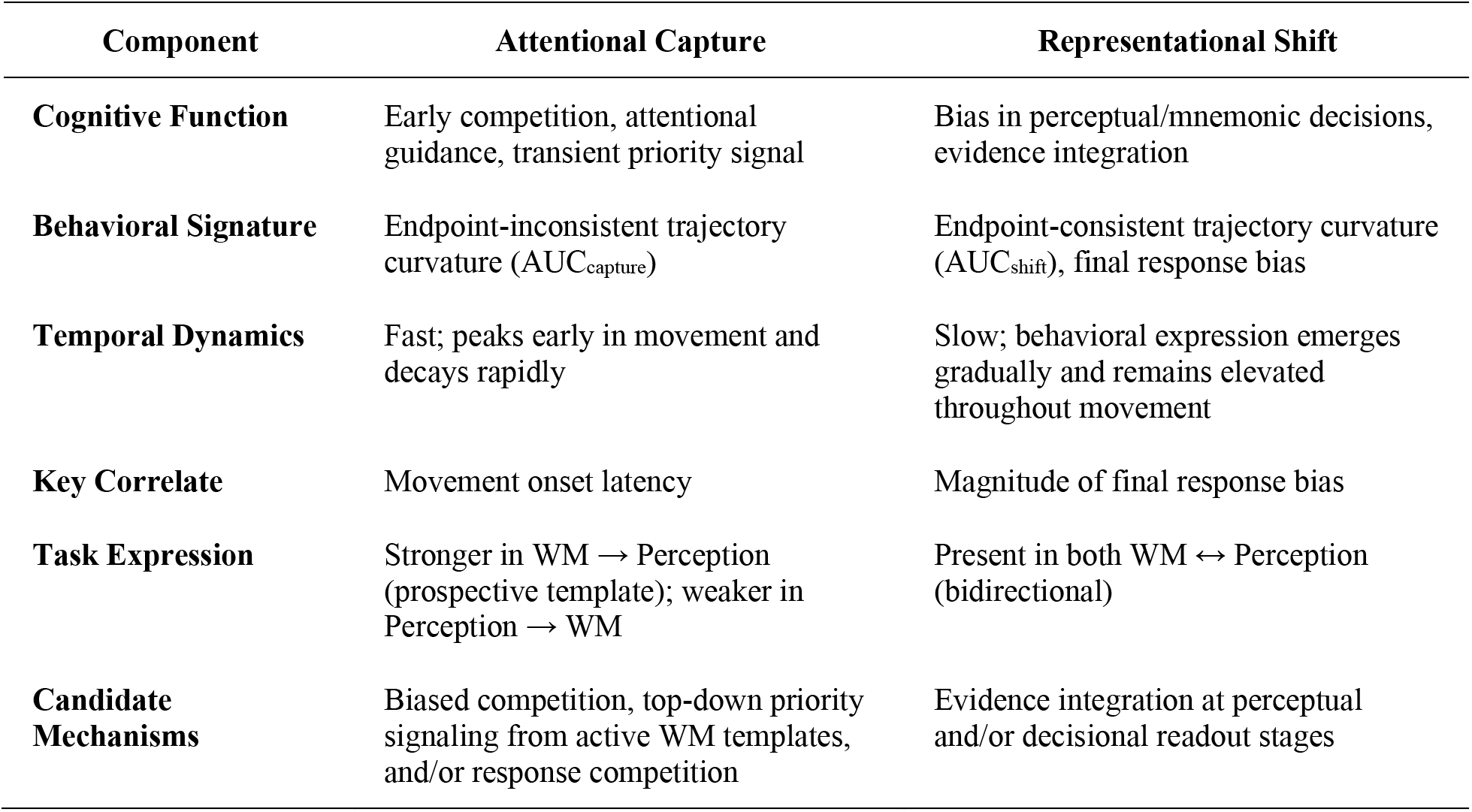
Distinct Temporal and Functional Components of Working Memory’s Top-Down Influence

| **Component** | **Attentional Capture** | **Representational Shift** |
| --- | --- | --- |
| **Cognitive Function** | Early competition, attentional guidance, transient priority signal | Bias in perceptual/mnemonic decisions, evidence integration |
| **Behavioral Signature** | Endpoint-inconsistent trajectory curvature ($AUC_{\text{capture}}$) | Endpoint-consistent trajectory curvature ($AUC_{\text{shift}}$), final response bias |
| **Temporal Dynamics** | Fast; peaks early in movement and decays rapidly | Slow; behavioral expression emerges gradually and remains elevated throughout movement |
| **Key Correlate** | Movement onset latency | Magnitude of final response bias |
| **Task Expression** | Stronger in WM → Perception (prospective template); weaker in Perception → WM | Present in both WM ↔ Perception (bidirectional) |
| **Candidate Mechanisms** | Biased competition, top-down priority signaling from active WM templates, and/or response competition | Evidence integration at perceptual and/or decisional readout stages |

Three converging findings support this conclusion. First, decomposing trajectories into endpoint-inconsistent and endpoint-consistent curvature revealed distinct relationships with movement onset latency and final estimation bias. The early, transient curvature (capture) was selectively correlated with how quickly a response was initiated but not with the final bias. In contrast, the endpoint-consistent curvature (shift) closely tracked the magnitude and direction of the final report, as expected given that it is defined relative to the endpoint path, but was not strongly related to response initiation speed. Second, time-resolved destination-vectors analyses confirmed distinct temporal profiles. The capture component peaked sharply near movement onset and quickly dissipated, whereas the behavioral expression of the shift component increased more gradually and remained elevated until the response was completed^1^. It is important to note that this temporal profile localizes when the endpoint-consistent bias became expressed in the unfolding movement, not when the underlying perceptual or mnemonic representation first became biased. Third, and most importantly, the expression of these two components was functionally distinct across directions of influence. When WM prospectively influenced perception, both capture and shift were evident. When perception reciprocally influenced WM, the shift remained strong while the capture-like component was markedly weaker. This asymmetry suggests that the relative expression of the two components depends on the direction and priority state of the interaction.

The fast-acting capture-like component is consistent with attentional guidance, but its motor expression does not uniquely identify an attentional locus. It may reflect a transient WM-driven priority signal, early competition between response tendencies, or both. This interpretation aligns with the biased competition model, which posits that active WM representations pre-activate corresponding feature-selective neural populations, granting a competitive advantage to matching sensory input for selection (Desimone & Duncan, 1995; Serences & Yantis, 2006). The initial deviation of the cursor toward the WM-matching color is consistent with an early priority signal or competing response tendency as the system resolves competition between the target and the memory-matching feature value. The transient rise and subsequent correction are consistent with a declining influence of the early competing tendency as task-relevant sensory evidence and the selected response increasingly constrain the movement. This pattern is also compatible with attentional disengagement and reorientation toward the primary task goal (Dieciuc et al., 2019). Time-resolved trajectory analysis thus provides a process-level window into how this early competition and subsequent correction unfold within a single decision.

The pattern observed in Experiment 2 further clarifies the conditions under which this early capture-like curvature is expressed. The capture-like component was strong when WM prospectively influenced perception and markedly weaker when the influence was from a just-encoded percept back onto memory (Park et al., 2017). This asymmetry suggests that the early capture-like influence depends on the priority state of the relevant representation rather than simply on the presence of mnemonic information when it is actively used to guide upcoming behavior (Park & Zhang, 2024). Neuroimaging studies support this distinction, showing that prospective WM templates engage frontoparietal networks that broadcast priority signals to sensory areas, whereas retrospective maintenance primarily involves sensory reinstatement without prioritization (Myers et al., 2017; van Loon et al., 2018). In this sense, a WM template that is actively guiding behavior is functionally distinct from a more passive trace of a previous percept, and the former is more likely to produce the rapid priority signal that manifests as transient motor capture toward memory-matching input.

The broader pattern of results is compatible with a precision-weighted integration framework, in which perceptual readout combines current sensory evidence with state-dependent priors supplied by WM. The similar magnitudes of the reciprocal endpoint biases in Experiment 2 argue against a purely task-specific artifact and instead support a general tendency for perceptual and mnemonic decisions to assimilate toward one another. The dissociation between the two trajectory components strengthens this interpretation, as early capture covaried with movement initiation latency but not endpoint bias, whereas sustained, endpoint-consistent curvature tracked endpoint bias independently of movement initiation latency. A single, undifferentiated decisional or motor account would predict shared variance across these measures and is therefore difficult to reconcile with this dissociation. The data instead point toward multiple stages at which WM can influence evolving decisions, from early competition between candidate responses to later integration of mnemonic and sensory information.

What do these findings imply about whether the locus of WM’s top-down influence is perceptual versus post-perceptual? A central challenge in debates over cognitive penetrability is distinguishing genuine perceptual changes from effects of attentional selection, decision rules, or response preparation (Firestone & Scholl, 2016). Our trajectory decomposition reveals temporally distinct movement signatures of an early capture-like influence and a sustained endpoint-consistent bias, and that the sustained, endpoint-consistent component is closely aligned with the final report. These observations are compatible with frameworks in which WM acts as a prior that shapes late perceptual readout and/or decision formation, but they do not by themselves rule out a decisional locus. From a broader theoretical perspective, the current results support a state-dependent view in which memory contributes flexible priors for perceptual judgments. In this view, WM belongs to a broader class of mnemonic influences that configure perception and decision making through dynamic, context-dependent processing of environmental regularities (Girshick et al., 2011), semantic knowledge (Hansen et al., 2006; Bannert & Bartels, 2013), and episodic retrieval (Bosch et al., 2014; Won et al., 2023).

The integrative and gradual nature of the sustained bias aligns with predictive coding and Bayesian integration frameworks of perception (Ma et al., 2006; Knill & Richards, 1996). In these frameworks, perception arises from the ongoing integration of sensory evidence with top-down expectations. WM representations can be viewed as prior beliefs about which features are most likely to occur in the current context. Such models combine these priors with incoming sensory evidence (the likelihood) to form an updated estimate (the posterior) that is consequently biased toward the prior. Within such accounts, gradual integration could naturally produce a more sustained bias than an early competitive influence. Within this framework, the present results are consistent with at least two temporally distinct contributions to perceptual decisions, one through an early competitive influence and another through a slower integrative bias, in which mnemonic content can contribute prior information to perceptual inference.

In conclusion, this study demonstrates that WM-related influences on perceptual decisions contain temporally distinct components. A rapid capture-like influence was strongest when a WM template was prospectively relevant, while a slower assimilative process was expressed in both directions of influence. Methodologically, by leveraging the temporal resolution of mouse-tracking and a variance-partitioning decomposition of trajectory deviations, the present approach offers a process-level view of decision dynamics that is sensitive to both transient and sustained influences within a single trial. This aligns with efforts to expand the rich behavioral repertoire for studying dynamic cognition (Nobre, 2022). More broadly, the findings support a view of cognition as a dynamic synthesis of past and present, with WM providing goal-dependent prior information that can shape how sensory evidence is weighted over time. This dynamic interplay allows perceptual decisions to balance stability with flexibility, supporting adaptive behavior in a changing environment (Kiyonaga et al., 2017).

## Ethics

This research was approved by the Institutional Review Board of the University of California, Riverside.

## Preregistration

### No aspects of the study were preregistered

^1^ Note that these temporal profiles are not a trivial consequence of the decomposition procedure. Although the shift component is defined with respect to the final response endpoint, this constraint only anchors the late portion of its destination-vector time series. Earlier values can in principle remain near zero, reverse in sign, or fluctuate. The fact that DVSHIFT tends to emerge gradually and remain elevated throughout the movement therefore reflects a genuine regularity in the trajectory dynamics, rather than a pattern that is mathematically inherited from the endpoint correction itself.

